# Agency as a Bridge to Form Associative Memories

**DOI:** 10.1101/2022.05.11.491543

**Authors:** Nicholas A. Ruiz, Sarah DuBrow, Vishnu P. Murty

**Author notes:** Correspondence concerning this article should be addressed to Nicholas A. Ruiz, Department of Psychology, Temple University, Weiss Hall, Room 532, 1701 N. 13th St, Philadelphia, PA 19122.

## Abstract

The perception of agency occurs when individuals feel their decisions exert control over their environment. While agency can increase memory for items, most real-life situations are more complex. The decisions we make not only affect the item we act upon, but all the other items in direct proximity of our decisions. Here, we examined how an individual’s agency to influence a situation affects their ability to learn associations between items that occur prior to and after making a decision. In our paradigm, participants were told they were playing a game show where they had to help a trial unique ‘contestant’ choose between three doors. On ‘agency’ trials, participants were allowed to pick any door they wanted. On ‘forced-choice’ trials, participants were instructed to select a door that was highlighted. They then saw the outcome, a ‘prize’ that was behind the selected door. Across two studies, participants show enhanced memory for contestants they saw in agency vs forced-choice trials. Memory benefits also extended to contestant - door and door - prize associations in both studies. Study 2 found this effect in the contestant - prize association. Notably, we found that agency also shaped the representation of memories such that they were stored as integrated sequences rather than individual relational pairs. Together, these data suggest agency over a situation leads to enhanced memory for all items in that situation. This enhanced binding for items may be occurring by the formation of causal links when an individual has agency over their learning environment.

---

Individuals are motivated to exert agency, such that they feel their choices and actions allow them to influence the external environment around them (Gallagher, 2012, Haggard, 2017, Haggard & Chambon, 2012, Moore, 2016). Studies on causal learning show that as early as infancy humans learn the relations between actions and the outcomes (Kuhn, 2012) and use this knowledge to act as a successful agent in their environment. Much of the existing literature on agency emphasizes the process of comparing the outcome of one’s own actions with their internal predictions of those actions (Haggard et al., 2002, Haggard, 2008, Wolpert et al., 1995) or generally how our action-outcome contingencies match our intentions (Chambon et al, 2014, Haggard, 2017). However, these processes have been limited in their characterizations of the effects of agency on episodic memory (Hon, 2017).

When an individual has agency over a choice, they experience a sequence in which they are cued to choose among a set of actions which subsequently dictate the outcome. Despite the majority of prior episodic memory literature focusing on outcomes, the association between cues, actions, and outcomes underlie the ability to guide future choice. How do individuals learn and remember the associations in a choice-sequence when we have agency? A key factor in exploring the intersection between memory and agency comes through the execution of a choice. Individuals’ choices bias representations of outcomes to be internally consistent in long-term memory (Mather et al., 2000), a mechanism theorized to be driven by cortico-striatal interactions (Delgado 2007, Leotti & Delgado, 2011, Leotti et al., 2010). Similarly, decision contexts modulate hippocampal memory via striatal and dopaminergic interaction to signal motivational significance (Shohamy & Adcock, 2010). Relevant to our current studies, motivationally-relevant studies can enhance the binding of cues with outcomes, however these studies have not manipulated individual’s agency in their decision environment (Rouhan & Niv, 2021). Yet, prior work suggests that many of these associative binding processes may actually reflect the act of making a choice, rather than reward motivation. For example, reward signaling in the striatum is present in choice situations but absent in non-choice situations (Tricomi et al., 2004). We propose that engagement of these motivated memory systems support memories for sequences of action-outcome contingencies and provide a scaffold to associate details within a decision sequence, when an individual has agency over the choice.

In line with these predictions, allowing individuals the opportunity to make a choice which influences their learning environment enhances item memory (Gureckis & Markant, 2012, Markant et al., 2016). If time is limited, individuals learn items more effectively when they choose what to study and those choices are honored (Kornell & Metcalfe, 2006). Control over learning environments has also been shown to improve learning and memory even when the choices being made are not directly related to the content of the to-be-learned items. For example, when given exploratory control over a learning environment, participants benefited from being able to control when a certain stimulus-location combination was presented (Voss 2011a). However, this mechanism was partially driven by the ability to revisit and re-study previously seen items (Voss 2011b). Prior work from our group has divorced active control and learning from stimulus timing, order, content, and presentation of the to-be-remembered items. In a series of studies, participants were given a choice to click on one of two ‘cards’ which would reveal a to-be-learned item. The cards were unrelated to the revealed items and did not control where or when the item would appear. Given this simple choice, Murty, DuBrow, and Davachi 2015, 2019 found participants better remembered items that appeared as a result of a participants choice compared to when they were forced to overturn one of the cards. While the existing literature shows how agency over a choice can positively affect memory for the outcome of a choice, it does not shed much light onto memory for the overall decision sequence, thus precluding the ability to understand how perceived choice influences associative memory processes more akin to event memory.

Given the role of motivated memory formation in associative binding, we hypothesize giving individuals agency over a similar decision sequence will boost associative memory for the elements embedded within a decision sequence. This idea is consistent with existing hippocampal theories of associative learning as well as its role in binding multiple elements of an experience (Eichenbaum et al., 2007, Mayes et al., 2007, Squire et al., 2004). In order to support adaptive memory formation, the binding of action to outcome only occurs when the action is voluntary and deliberate, as these represent states when internal predictions match with sensory outcomes (Ebert & Wegner, 2010, Frith, 2014, Haggard et al., 2002, Moore & Obhi, 2012). However, little work has been done evaluating how this binding may be expanded to the domain of memory. Work that has explored this effect either test memory for the item that is acted upon (Hon & Yeo, 2021) or the outcome of the action (Murty et al., 2015, 2019) but not for the associations between items.

In the current study, we aim to explore how agency over a choice-sequence will affect associative memory for the components of the sequence, and further delineate the nature of this underlying representation. Participants are told they are participating in a game show, where they will have to help a trial-unique contestant choose between one of three doors. Behind each door will be a prize, and they must help the contestant win the best prize possible. Critically, participants will either get to choose any of the three doors freely, “agency” trials, or they will be forced to select a door by the experimenter, “forced-choice” trials. We expect agency to act as a bridge, bolstering associative memory for: items that occur prior to a choice, the choice item, and the item that appears as a result of the choice. Further, we ran a series of post-hoc analyses to better understand the underlying nature of associative memory representations, delineating whether they are stored as contingent sequences or independent relationships.

## Method

### Participants and Design

179 participants were recruited across two studies (study 1 n = 48; study 2 n = 131) via Prolific.ac, an online subject pool for behavioral studies (Palan & Schitter, 2018). To qualify for this study, participants must have reported being 18-35 years old, having normal or corrected-to-normal vision, living within the United States, and speaking fluent English. Participants were required to use a desktop or laptop computer, use of mobile devices was restricted. Sample sizes for both studies were determined a priori using a power analysis (Cohen, 1988) with alpha = 0.05 and power = 0.95. We based the sample size of study 1 on an effect size from a previous study (Murty, DuBrow, & Davachi, 2015) which required participants to remember an item based on a choice without testing associative memory. The sample size from study 2 was based on effect sizes obtained from study 1, however this yielded a much larger sample size suggesting that study 1 may have been slightly under-powered. Participants’ data were excluded from the final analysis if they did not respond to at least 75% of the encoding trials, 75% of the retrieval trials, or made repetitive responses on 50% or more consecutive trials for retrieval phase 2 or retrieval phase 3 (see below). This resulted in an n = 43 for study 1 and an n = 111 for study 2. Participants were paid at a rate of $8-$10 / hour for their participation.

### Materials and Procedure

Informed consent and stimuli were presented using Inquisit, an online-based experiment hosting website (Grootswagers, 2020). Study 1 and Study 2 contained the same stimuli and procedures. Study 2 was a pre-registered, replication of study 1 (AsPredicted #58695). The experimental stimuli consisted of cartoon figures generated within the Toca Boca application (https:tocaboca.com/apps), cartoon doors with patterns, and photographs of neutral, man-made objects (e.g. blender). The Toca Boca characters were used in a previous study and all contained neutral facial expressions (Murty et al., 2020). 7 cartoon-patterned doors were created using royalty-free templates from www.vecteezy.com. These doors were pre-screened in a separate group of participants (n = 43) who rated their likeability on a scale from 1-5. The three highest rated doors (water pattern [*m* = 3.23, *sd* = 1.43], leaf pattern [*m* = 3.62, *sd* = 1.26], and lego-brick pattern [*m* = 2.85, *sd* = 1.40]) were chosen for the experiment. 197 object images were also pre-screened by the same group of subjects who rated whether they knew what the object was as well as how much they would like to win that object on a game show on a scale from 1-5. From this, 120 objects were selected based on at least 80% of the subjects having reported knowing what the object was. From these 120 objects, stimulus lists containing 60 Toca Boca characters and 60 objects were created for both studies, with each list counterbalanced by preference for the items on that list. For study 1, six stim-lists were created and presented to groups of 8 participants. For study 2, five stim-lists were created and presented to groups of 25-27 participants. All participants completed an incidental encoding gameshow task, a short working memory task, and a retrieval task. Results from the working memory task were not analyzed and will not be reported here, the purpose of this task was to provide filler (average completion time: 5 minutes) between the encoding and retrieval tasks.

The first segment of the task was the incidental encoding task. The incidental encoding task (**Fig. 1**) consisted of 40 trials and was modeled after a gameshow. On each trial, participants were told they were in a game show and they had to assist a trial unique “contestant” (Toca Boca character) in choosing between one of three doors, each of which were numbered. The three doors remained in the same position for each trial, each subject, and across both studies so spatial location and door identity could not be dissociated. The position of the doors was randomly selected. Participants were told that behind each door is a prize, but some doors lead to better prizes. This was to encourage them to fully explore each door. Their task was to help each contestant win the best prize possible. Unbeknownst to participants, the object image presented was pre-determined by the experimenter in a randomized manner, thus allowing us to manipulate perceived agency rather than actual control over learning. Each trial in the encoding task consisted of a sequence of images. First, participants saw the contestant and the three doors on the screen. They were given 10s to choose one of the three doors for the contestant. After a choice was made, participants saw the contestant and the chosen door (2s), and finally they saw the contestant and the unique prize for that trial (2s). There was a 2s inter-trial interval and then the next trial began.

**Figure 1.**
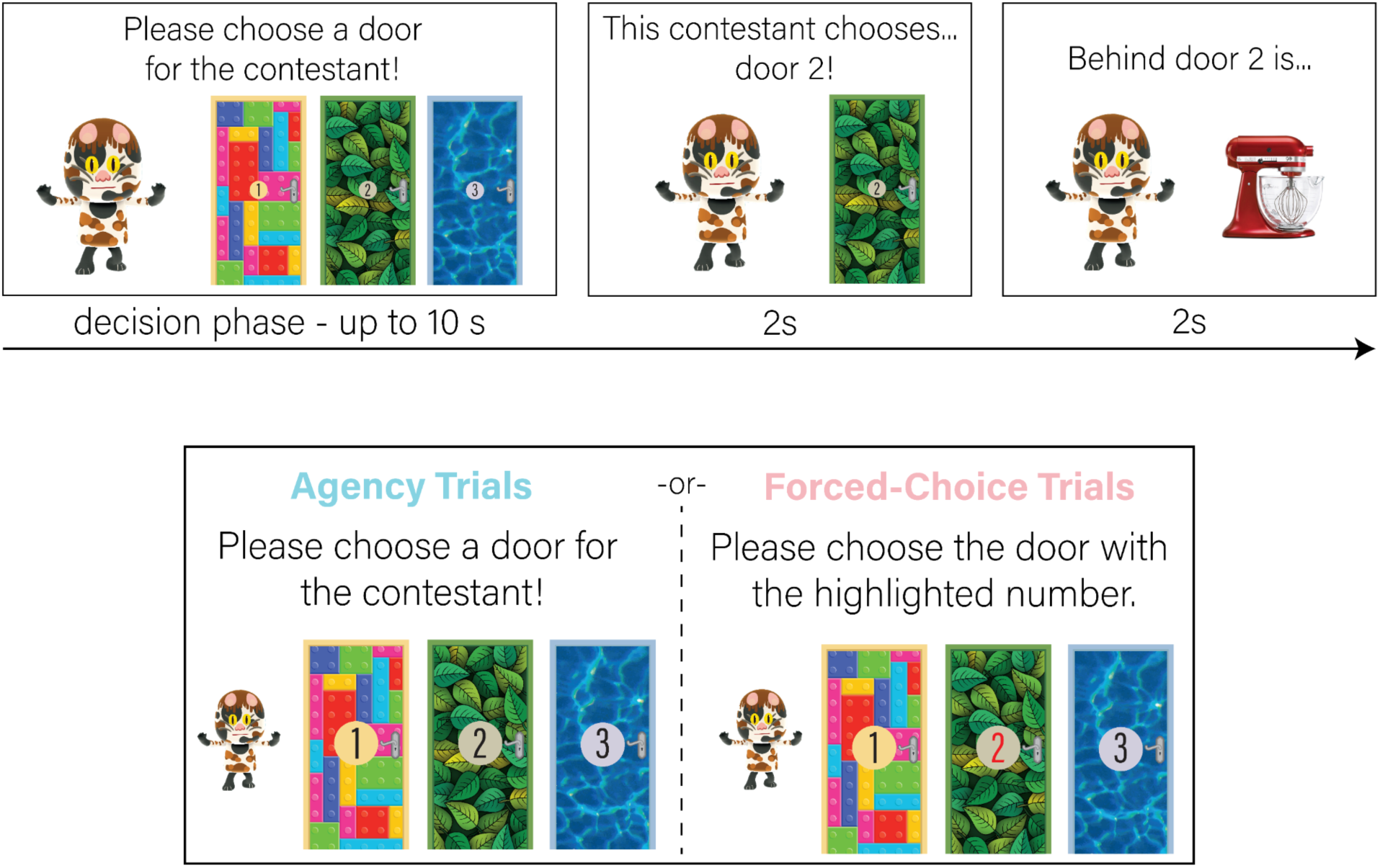
Incidental Encoding Phase. Participants were told they were participating in a game show, where they had to help trial-unique contestants select a door. They were told that behind each door was a different type of prize and some doors contained better prizes than others despite object image pairings being pre-determined by the experimenter. At the start of each trial, participants saw the contestant and the three doors. They were either asked to choose any door for the contestant (agency trials) or select the door with the highlighted number (forced-choice trials). They were given up to 10 s to make a choice. After choosing one of the three doors, they viewed the contestant and the door that was selected. Finally, they saw the contestant and the prize that was behind the selected door.

There were two types of trials: agency and forced-choice (**Fig. 1**). On agency trials, participants were informed they were allowed to choose any of the three doors they wanted. On forced-choice trials, participants were instructed to choose the door that had a highlighted number (the number was changed to red). Participants would make a choice by clicking on the door itself. There were 20 agency trials and 20 forced-choice trials, across 8 blocks consisting of 5 trials each. Each block alternated between agency and forced-choice trials, with the trial type of the first block being counterbalanced across participants.

After the encoding task, participants completed a short working memory task, the visual digit span. This task is provided by Inquisit based on Wood et al., 2011. Data from this task was not analyzed and served only as a filler between the encoding and retrieval tasks.

Finally in the last segment after the working memory task, participants completed a surprise retrieval task which consisted of three sub-phases (**Fig. 2**). The first phase consisted of 60 trials, on each trial participants were presented with either a contestant from the encoding phase or a novel contestant (40 old contestants, 20 new contestants; **Fig. 2a**). They were instructed to indicate whether they remembered the contestant from the encoding task (yes) or not (no). If they responded yes, they were shown three objects from the encoding phase. They were asked to indicate which prize that contestant won. One of the prizes was the correct prize associated with that contestant and the other two prizes were seen in the encoding phase associated with different contestants. The second retrieval test consisted of 40 trials in study 1 and 60 trials in study 2 (**Fig. 2b**). Participants were again presented with either old or new contestants and were instructed to indicate which door that contestant chose. The three doors from the encoding phases were presented in the same order as they were during encoding. Study 1 only contained 40 trials due to a programming error, therefore not all old contestants were seen in this phase for participants in study 1. On average, in study 1 participants saw 26.9 old contestants and 13.1 new contestants during this phase. All analyses reflect this difference in presentation number. The third phase of retrieval consisted of 40 trials (40 old prizes; **Fig. 2c**). Participants were presented with each of the 40 prizes from the encoding task and were instructed to indicate which door that prize was behind. Each of the retrieval tasks were self-paced, with a maximum response time of 10 s. Finally, participants completed a post-task questionnaire and were paid for their time.

**Figure 2.**
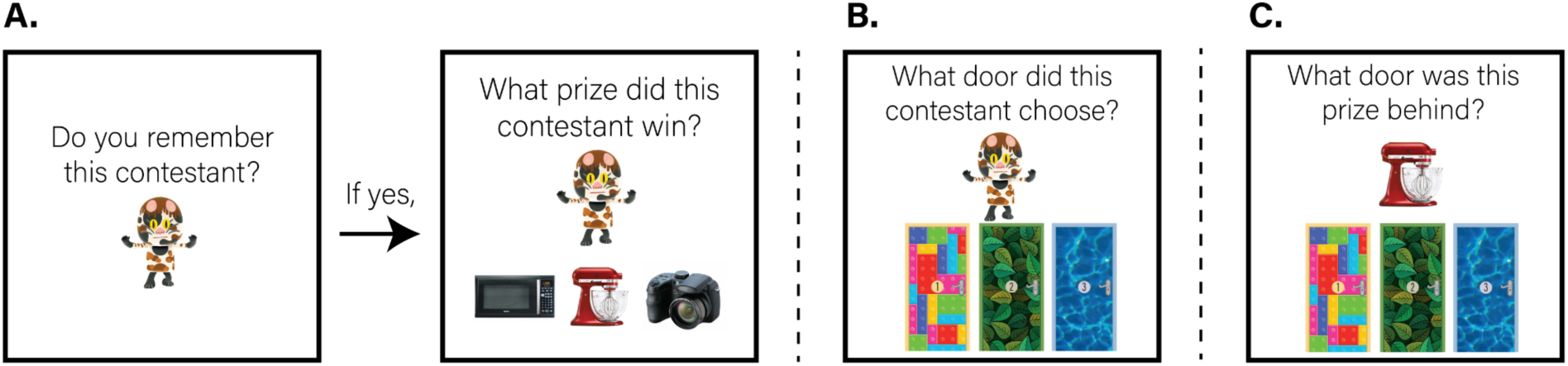
Retrieval Phases. **(A.)** Contestant recognition and contestant - prize associative memory tests. In a surprise retrieval test, participants were told they would see various characters and were to indicate by making a “yes” or “no” response as to whether they remember seeing that character as a contestant in the game show task. If they responded “yes”, they would then see the same character and three prizes. They were then asked to indicate which prize that character won. The three prizes had all previously been viewed in the encoding phase. **(B.)** Contestant - door associative memory test. Participants were told they would again view various characters who may or may not have been in the game show task. They were instructed to select the door that the character chose during the encoding phase. **(C.)** Prize - door associative memory test. Participants were told they were going to see all of the prizes from the game show task and had to indicate which door that prize was behind. Participants saw the three retrieval phases in the order presented in this figure.

### Statistical Analysis

To examine deliberation differences during the choice phase of the encoding task, reaction time was first analyzed on a between-subjects level. Mean reaction times during the choice phase for each condition (agency and forced-choice) were calculated and compared using t-tests. This analysis was conducted across the two studies. For all analyses using t-tests, reported p values are two-tailed, with values less than .05 considered statistically significant. 95% confidence intervals are reported where appropriate. Effect sizes (cohen d for t-tests, eta-squared for ANOVA) were calculated using the ‘effectsize’ and ‘lsr’ packages in R (v 0.4.5).

Below, we describe the summary statistics we generated for each individual participant for each of our three retrieval tasks. For contestant recognition memory, we first determined whether participants performed above chance levels of performance by comparing hit rates and false alarm rates. Then we compared corrected recognition (hit rate minus the false alarm rate) of contestant memory across conditions. Contestant - prize, contestant - door, and door - prize memory was calculated using accuracy. For all analyses, data was compared across conditions using paired t-tests.

Next we ran a series of control analyses. First, we examined if differences in reaction times across conditions influenced subsequent memory. To do this, we used generalized linear mixed effects models to examine the relationship response times and memory outcome. These models were implemented using the lme4 package in R (lme4 v 1.1-26; R v 4.0.3), using a model comparison approach where we determined how the addition of another factor influenced the overall model fit for each of our four memory tests. As we predict condition (agency, forced-choice) will be the strongest predictor of memory outcome, we first created a baseline model which predicts memory outcome by condition. This baseline model was compared to a reaction time model which predicts memory outcome by condition and reaction time (during the choice phase). Finally, the reaction time model was compared to an interaction model which predicted memory outcome by condition, reaction time, and the interaction between condition and reaction time. All models were computed on a trial-wise level and included ‘subject’ as a random effect to account for within-subjects variation in the data. We used analysis of variance (ANOVA) to conduct model comparisons. Data from both studies were combined for this analysis.

We next were interested in whether a preference bias during the choice phase may have influenced memory outcome performance. Choice preference across the two studies was assessed using a chi square test. Specifically, we compared participants’ idiosyncratic preferences in selecting each of the three doors against 1/3rd for each door. Next, we examined if any subject-level bias had an influence on memory performance. If a participant preferred one of the three doors, and therefore had a response bias towards that door during the agency choice phase, this would cause an inflated baseline or ‘chance’ level performance during retrieval phases two (contestant - door memory) and three (prize - door memory). To correct for this in all the analyses detailed below, we conducted a permutation analysis where we shuffled responses on the relevant tests for each participant 10,000x to generate a subject-specific idiosyncratic null distribution. We then took the average of these permutations and subtracted it from each subject’s actual performance. In this way, for these two measures, scores represent an unbiased retrieval score as they separate individual subject biases from their actual performance. We then conducted t-tests on the difference scores (actual performance minus permutation performance) to compare performance across conditions. Data from both studies was combined for this analysis.

Finally, we were interested in examining whether having agency over the decision sequence was leading participants to view the items as a sequence of interconnected events rather than sets of individual pairs (see **Fig. 4a-c**). To probe this, we examined how performance on contestant (A) - door (B) and door (B) - prize (C) corresponded to performance on contestant (A) - prize (C). While the A-B (contestant - door) and A-C (contestant - prize) pairs were seen together during the encoding phase, the B-C (door - prize) pair was never seen together. If participants viewed the sequence as individual items of A, B, and C, successful retrieval of the A-C pair should be contingent on successfully retrieving the first item A, its relationship with item B, then recalling B’s relationship with item C. Whereas remembering the pairs as unitize, relational pairs would indicate participants could successfully recall the A-C pair without needing to recall A-B or B-C. We examined A-C performance with a 2 (condition: agency, forced choice) x 2 (A-B + B-C performance: 1 if both were correct [sequence intact], 0 otherwise [sequence fragmented]) analysis of variance, testing for main effects of condition, A-B + B-C performance, and their interaction. For this analysis, we collapsed data across studies to increase statistical power.

**Figure 3.**
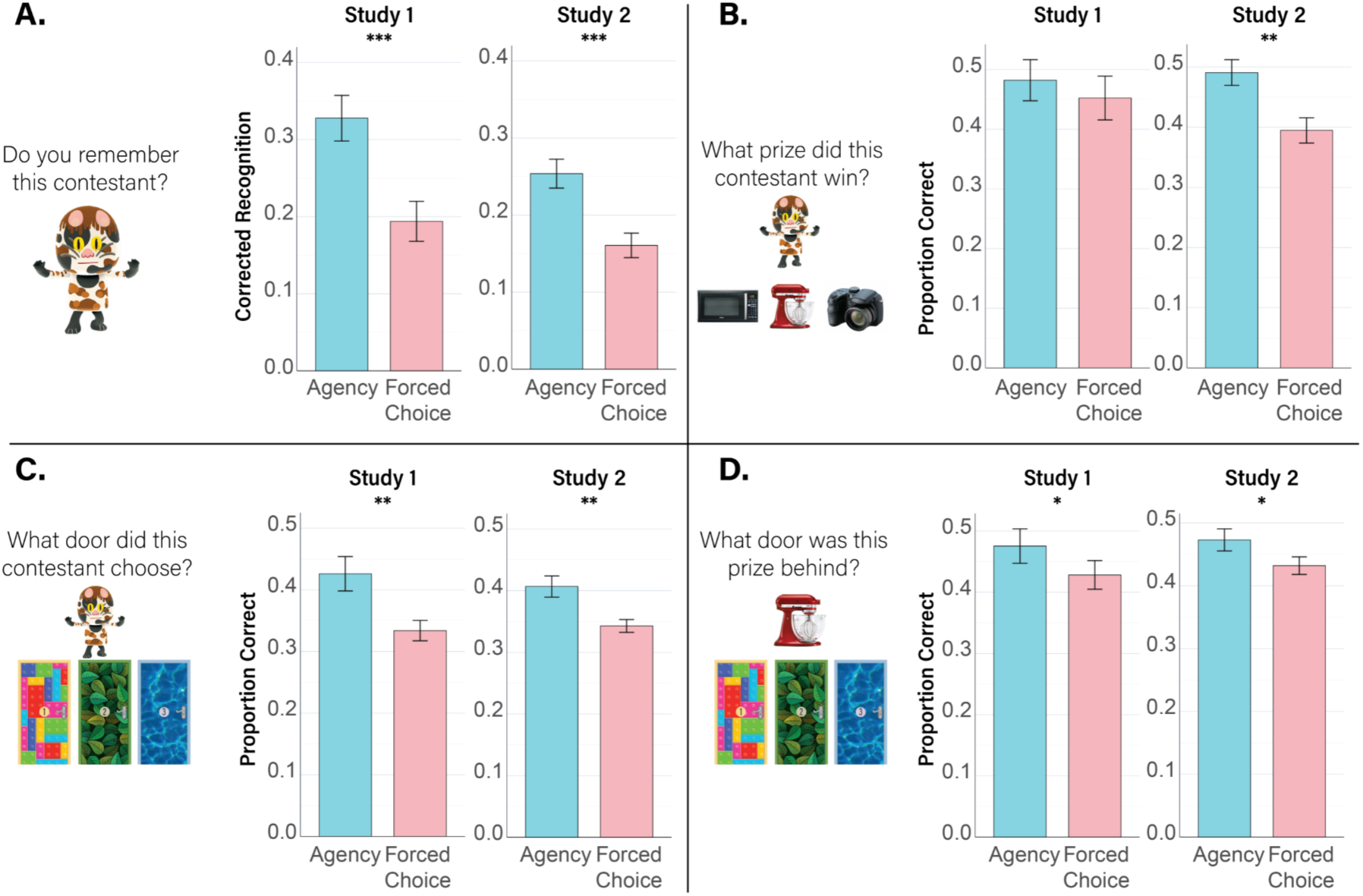
Retrieval Results. **(A.)** Contestant corrected recognition. Across both studies, there was a significant difference in corrected recognition between contestants viewed in agency trials vs. contestants viewed in forced-choice trials. **(B.)** Contestant - prize associative memory accuracy. For study 1, there was no statistically significant difference across contestant - prize associative pairs that occurred in agency trials vs. forced-choice trials. For study 2, contestant - prize associative pairs that occurred in agency trials had statistically higher accuracy than pairs that occurred in forced-choice trials. **(C.)** Contestant - door associative memory accuracy. Across both studies, contestant - door associative pairs that occurred in agency trials had significantly higher accuracy than pairs that occurred in forced-choice trials. **(D.)** Prize - door associative memory accuracy. Across both studies, prize - door associative pairs that occurred in agency trials had significantly higher accuracy than those that occurred in forced-choice trials. Error bars indicate *SEM*. \**p < 0*.*05 ** p < 0*.*01 *** p < 0*.*001*.

**Figure 4.**
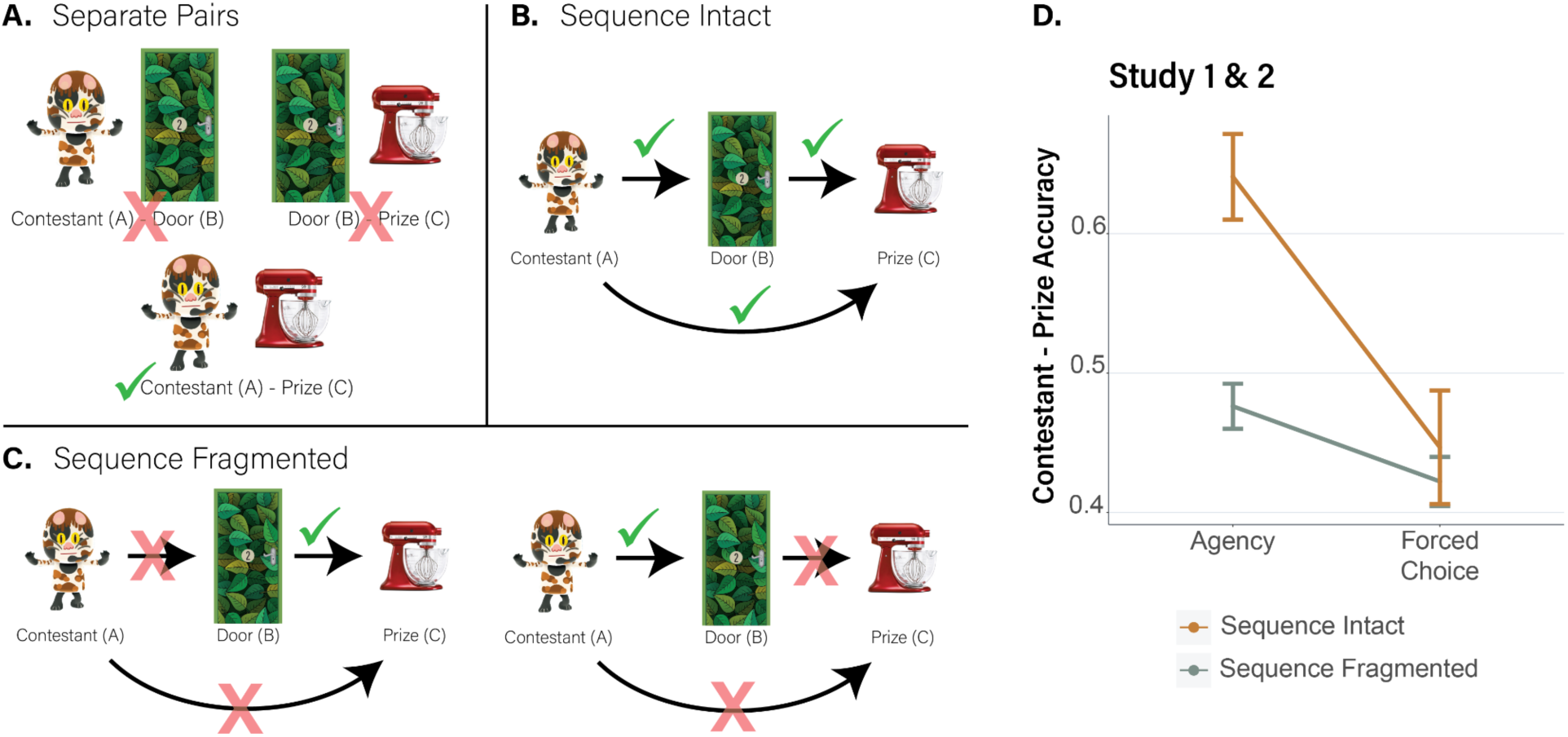
Sequence Conceptualization and Contingency Analysis. **(A.)** Separate Pairs. During encoding, participants viewed pairs of items on the screen at the same time. The contestant (A) - door (B) pair was first seen, then the contestant (A) - prize (C) pair were seen. The door (B) - prize (C) pair were never seen on the screen together. If participants viewed these item pairs as a singular item, recall of pairs further in the sequence should not rely on pairs that occurred earlier in the sequence. For example, one could still recall the A-C pair if they viewed it as a singular pair, independent of the previous pairs A-B and B-C. **(B.)** Sequence Intact. If individuals viewed the items in the sequence as individual items the recall of the A-C pairing would be contingent on recalling the earlier pairs A-B and B-C. **(C.)** Sequence Fragmented. If one did not recall either the A-B or the B-C pair, the sequence would be fragmented such that they would not be able to recall the A-C pair. **(D.)** Participants were significantly more likely to recall the contestant (A) - prize (C) associative pair if they also successfully recalled the both contestant (A) door (B) and door (B) - prize (C) pairs. The gold line (sequence intact) indicates both the contestant - door and door - prize pairs were successfully recalled. The gray line (sequence fragmented) indicates either one or both of the pairs were not successfully recalled. This effect was significantly larger for pairs that occurred in agency trials vs. forced-choice trials.

### Data Availability

Stimuli, experimental code, analysis code and data can be found at: https://osf.io/ek53n/.

## Results

### Analysis of Choice Phase

We first examined whether agency leads to differences in reaction times during the choice phase. Across both studies, reaction times were faster in the forced-choice condition versus the agency condition (Study 1: forced-choice, mean = 2,173 ms, *sd* = 696 ms; agency, mean = 2,642 ms, *sd* = 927 ms; t(42) = 3.60, p < 0.001; Study 2: forced-choice, mean = 2,314 ms, *sd* = 920 ms; agency, mean = 2,616 ms, *sd* = 1,130; t(110) = 3.43, p < 0.001).

We next examined whether there was a response bias in the choice phase at the group-level and if any bias could partially explain any memory difference. Our analysis first examined the frequency of which each of the three doors was chosen across all participants, where we found no significant difference in the number of times a door was chosen compared to chance in either study (study 1: χ^2^(4) = 1.16, p = 0.88; study 2: χ^2^(4) = 1.29, p = 0.86).

### Contestant Memory

Memory for each contestant was probed in the first phase of retrieval (**Fig. 2a**) using corrected recognition (hits minus false alarms; see **Table 1** for means and standard deviations). Memory was above chance in both conditions for both study 1 (t’s (42) > 8.01, p’s < 0.001) and study 2 (t’s (110) > 10.20, p’s < 0.001). We next compared corrected recognition (hit rate minus false alarm rate) across conditions (**Fig. 3a**). Across studies, corrected recognition was enhanced for contestants from the agency condition versus those in the forced-choice condition (Study 1: t(42) = 4.73, p < 0.001, d = 0.78; Study 2: t(110) = 5.95, p < 0.001, d = 0.49).

**Table 1.**
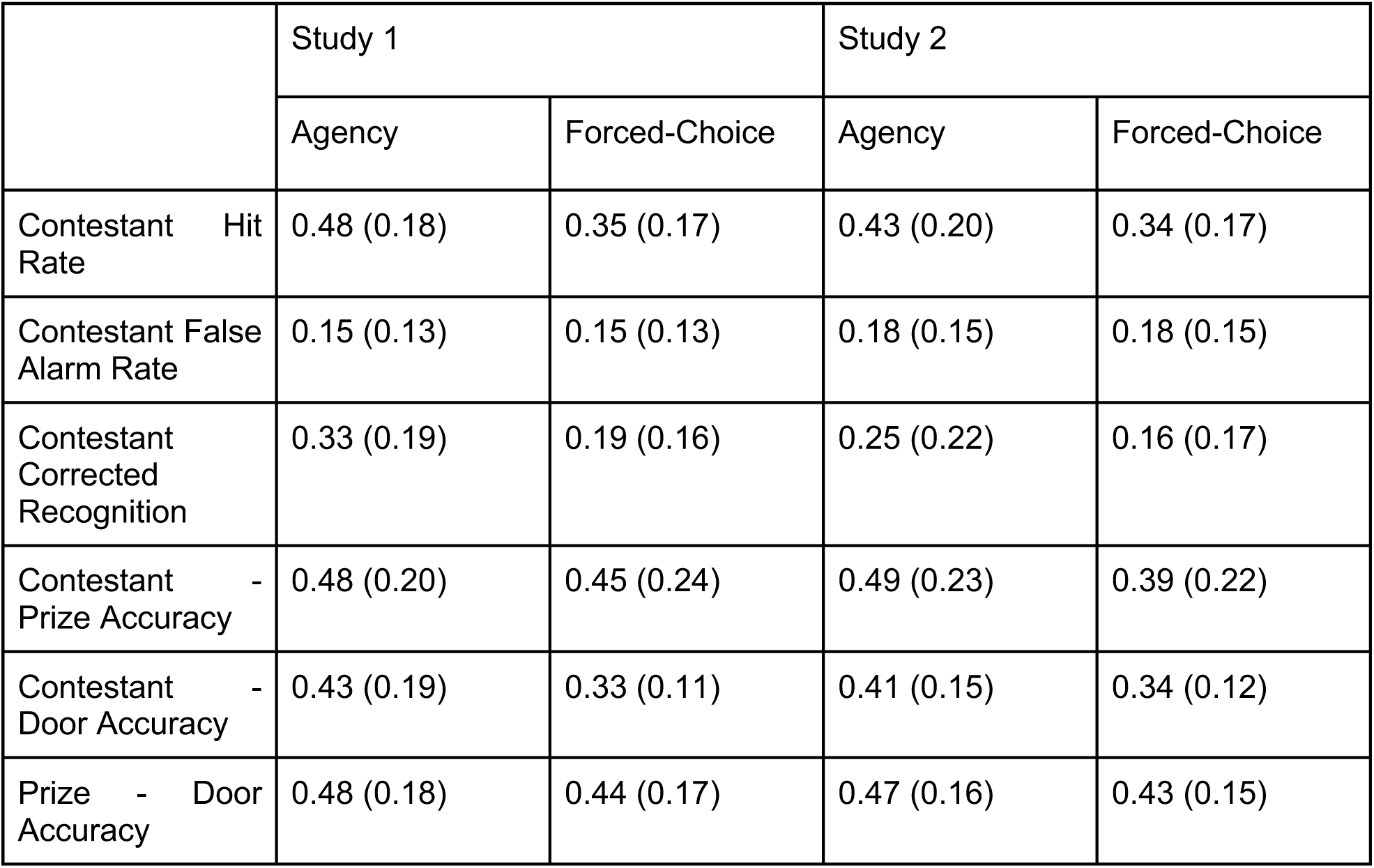
Mean (standard deviation) scores for all memory tests across agency and forced-choice trials for Study 1 and Study 2.

### Associative Memory

As part of phase 1, contestant - prize associative memory (accuracy) was probed if the participants reported remembering the contestant (**Fig. 2a**). Memory for contestant - prize pairs was above chance (chance = 1/3rd) for both agency and forced-choice pairs in study 1 (agency: t(40) = 4.87, p < 0.001; forced-choice: t(41) = 3.18, p = 0.003) and in study 2 (agency: t(110) = 7.28, p < 0.001; forced-choice: t(109) = 2.94, p = 0.004). In study 1, there was no difference in memory for contestant - prize associative memory for pairs associated in the agency condition versus pairs associated in the forced-choice condition (t(40) = 1.01, p = 0.32, d = 0.13; **Fig. 3b**), albeit numerically memory was greater in the agency condition. With increased sample size, study 2 showed enhanced memory for contestant - prize pairs when those pairs occurred in the agency condition versus pairs from the forced-choice condition (t(109) = 3.45, p < 0.001, d = 0.43; **Fig. 3b**).

In the second phase of retrieval, contestant - door associative memory (accuracy) was probed (**Fig. 2b**). Memory for contestant - door pairs was above chance (chance = 1/3rd) for agency pairs (t(42) = 3.21, p = 0.003) but not forced-choice pairs (t(42) = 0.03, p = 0.97) in study 1; and in study 2 (agency: t(110) = 5.00, p < 0.001; forced-choice: t(110) = 0.86, p = 0.39). In study 1, participants better remembered the contestant - door associative pair if the pair occurred in the agency versus forced choice condition t(42) = 2.86, p = 0.007, d = 0.59 (**Fig. 3c**). This result was replicated in study 2 where memory for contestant - door associative pairs from the agency condition were enhanced compared to pairs in the forced choice condition (t(110) = 3.68, p < 0.001, d = 0.47; **Fig. 3c**). We next ran a permutation analysis to control for the influence of any subject-level response bias during the choice phase on memory. After subtracting the mean of the permutations from the actual performance of any given subject, a paired t-test revealed memory for the contestant - door association was enhanced for pairs from the agency condition versus the forced-choice condition (t(153) = 4.92, p < 0.001, d = 0.56).

The final phase of retrieval probed associative memory for door - prize associations (**Fig. 2c**). Memory for door - prize pairs was above chance (chance = 1/3rd) for both agency and forced-choice pairs in study 1 (agency: t(42) = 5.28, p < 0.001; forced-choice: t(42) = 3.57, p < 0.001) and in study 2 (agency: t(110) = 9.42, p < 0.001; forced-choice: t(110) = 7.07, p < 0.001). For study 1, participants better remembered pairs from the agency condition versus pairs from the forced-choice condition t(42) = 2.18, p = 0.03, d = 0.27 (**Fig. 3c**). This result was replicated in study 2, where memory for agency pairs was enhanced compared to forced-choice pairs (t(110) = 2.59, p = 0.01, d = 0.27; **Fig. 3d**). A permutation analysis was conducted to explore whether a subject-level response bias during the choice phase influenced memory between conditions. After subtracting the mean of the permutations from the actual performance of any given subject, a paired t-test revealed memory for the door - prize association was enhanced for pairs from the agency condition versus the forced-choice condition (t(153) = 3.51, p = 0.001, d = 0.28).

### Control Analysis for Reaction Time Differences Across Conditions

The above results show enhanced items and associative memory when the participants had agency over the decision sequence vs when they were forced to choose. However, the memory results may have been confounded with differences in reaction time occurring during encoding. To determine if differences in memory across condition results from reaction time, we characterized how predictive reaction time and condition were of memory outcome for contestant recognition, and associative memory.

First, we compared generalized linear mixed effects models examining the role of reaction time (RT) during the choice phase and condition on contestant recognition. Although including reaction time during the choice phase into the baseline model significantly increased the model fit (model comparison: χ^2^(1) = 8.28, p = 0.004; baseline model: AIC = 7808.6, BIC = 7828.8; RT model: AIC = 7802.3, BIC = 7829.2), condition remained a significant predictor of recognition memory (β(6101) = -0.47, p < 0.001, 95% CI = [-0.58, -0.36], SE = 0.06, z = -8.41). This finding suggests that while encoding reaction time may significantly predict recognition memory, it did not undermine the influence of agency on recognition memory. Adding a condition*RT interaction did not increase the model fit (model comparison: χ^2^(1) = 0.38, p = 0.54; interaction model: AIC = 7804.0, BIC = 7829.2; reaction time model: AIC = 7802.3, BIC = 7829.2), suggesting that reaction time could not account for agency-related memory benefits. Finally, to examine whether condition would improve the fit, we compared a model which includes both RT and condition to a model that only includes RT. The model which included RT and condition had a significantly better model fit compared to the RT alone model (model comparison: χ^2^(1) = 71.12, p < 0.001; RT + condition model: AIC = 7802.3, BIC = 7829.2; RT alone model: AIC = 7871.5, BIC = 7891.6).

When examining contestant - prize associative memory, adding RT significantly increased the model fit (χ^2^(1) = 14.78, p = 0.0001; baseline model: AIC = 3271.8, BIC = 3289.1; reaction time model: AIC = 3259.0, BIC = 3282.1), however condition remained a significant predictor of associative memory (β(2386) = -0.30, p < 0.001, 95% CI = [-0.47, -0.13], SE = 0.09, z = -3.49). This again suggests that while encoding reaction time may significantly predict contestant - prize memory, it did not undermine the influence of agency. Adding a condition*RT interactions (χ^2^(1) = 1.94, p = 0.16; interaction model: AIC = 3259.1, BIC = 3288.0; reaction time model: AIC = 3259.0, BIC = 3282.1) did not improve the model fit. To again examine whether condition would improve the fit, we conducted a comparison of an RT + condition model to a model with only RT. The RT + condition model indeed had a significantly better model fit than the RT alone model (model comparison: χ^2^(1) = 12.22, p < 0.001; RT + condition model: AIC = 3269.3, BIC = 3286.6; RT alone model: AIC = 3259.0, BIC = 3282.1)

When examining contestant - door associative memory, adding RT did not significantly increase the model fit (χ^2^(1) = 3.27, p = 0.07; baseline model: AIC = 7205.1, BIC = 7224.9; reaction time model: AIC = 7203.9, BIC = 7230.3) and condition was the only significant predictor of contestant - door associative memory (β(5430) = -0.34, p < 0.001, 95% CI = [-0.45, -0.23], SE = 0.06, z = -6.01). Adding a condition*RT interaction did not significantly increase the model fit (χ^2^(1) = 2.03, p = 0.15; interaction model: AIC = 7203.8, BIC = 7236.8; reaction time model: AIC = 7203.9, BIC = 7230.3).

Finally, when examining prize - door associative memory, adding RT did not increase the model fit (χ^2^(1) = 2.63, p = 0.10; baseline model: AIC = 8144.9, BIC = 8165.0; reaction time model: AIC = 8144.3, BIC = 8171.1) and condition was the only significant predictor of prize - door memory (β(5986) = -0.21, p < 0.001, 95% CI = [-0.31, -0.10], SE = 0.05, z = -3.92). Adding a condition*RT interaction into the model did not increase the model fit compared to the reaction time model (χ^2^(1) = 0.04, p = 0.84; baseline model: AIC = 8146.2, BIC = 8179.7; reaction time model: AIC = 8144.3, BIC = 8171.1).

### Analysis of the Underlying Representation of Associative Memories

The results thus far suggest that agency increases pairwise associative memory. However, these prior analyses do not discriminate whether agency enhances memory separately for each individual pair, or rather whether agency facilitates the binding of all the associations into one integrated sequence. If agency facilitates memory integration, there should be inter-dependence among memory measures, such that successful contestant - prize (A-C) memory would depend on successfully recalling the entire sequence (sequence intact), both contestant - door (A-B) and door - prize memory (B-C). If either one or both of the constant - door (A-B) or door - prize (B-C) were not successfully recalled (sequence fragmented), then one would be unable to recall the contestant - prize (A-C) pair. To examine whether contestant (A) - prize (C) memory was contingent on remembering the other pairs of the decision sequence (contestant (A) door (B), door (B) - prize(C)), we conducted a 2 (condition: agency, forced choice) x 2 (sequence memory: 1 if intact, 0 if fragmented) analysis of variance on A-C memory (**Fig. 4d**). We found a significant main effect of condition, *F*(1,2139) = 14.92, p < 0.001, η^2^ = 0.006) such pairs seen in the agency condition were more likely to be recalled than those that appeared in the forced condition. We also found a significant main effect of sequence outcome, *F*(1,2139) = 15.75, p < 0.001, η^2^ = 0.007, such that, when the sequence was intact (both the A-B and B-C pairs were recalled), participants were more likely to recall the A-C pair. Critically, we found a significant condition by sequence outcome interaction, *F*(1,2139) = 6.13, p = 0.01, η^2^ = 0.003, such that memory for the A-C (contestant - prize) pair was contingent on correctly recalling both the A-B (contestant - door) pair as well as the B-C (door - prize) pair. This effect was particularly higher when the participants had agency over the sequence compared to when they did not (forced trials), suggesting that agency increases the contingency of individual elements of the decision sequence.

## Discussion

The sense of agency refers to the feeling when one’s own choices can exert control on their environment and is a key factor in many aspects of the human experience. In the current study we were interested in examining how giving individuals’ agency to make a choice within a decision sequence could influence associative memory for all the items embedded in their decision sequence. Here we provide evidence that the sense of agency enhances memory for items associated with a choice, and facilitates the formation of associations between items, choices, and outcomes. Further, we show evidence that agency may enhance binding such that the items within the associations are not represented as individual pairs but as an integrated sequence where recall of the outcome is contingent on recall of the intervening items. Together we show that imbuing individuals with agency during learning with a choice benefits memory for items proximal to that choice and facilitates the binding of the items into an integrated sequence.

We have shown a simple manipulation in agency can enhance memory for items and item pair associations in a sequence. Giving individuals agency over what information to learn and how much time to spend studying has been shown to enhance learning of facts (Metcalfe 2002, Metcalfe & Kornell 2003, 2005). However, the learning effects seen in these studies are in part due to metacognitively guided decisions to select easier information to allow for more learning in a given amount of time (Kornell & Metcalfe 2006). The choices the participants made in these studies are guided by higher order processes, involving probing one’s own knowledge base to optimally select information that has a higher probability of being retained given time constraints. These stand in stark contrast to the relatively simple choices participants made between doors in the current study, where either choice would have no control over difficulty or timing of stimuli presentation. Complementary work allowing individuals to actively sample information about novel categories has shown learning enhancements when the choices being made are self-directed (Markant & Gureckis, 2010, Gureckis & Markant, 2012). The authors liken this effect to a child pointing to an unfamiliar animal in a book and asking for its name rather than waiting for a parent to give this information, a self-directed process thought to be motivated by reducing uncertainty (Markant & Gureckis 2012a, 2012b). While these studies show the benefit of having control over certain aspects of one’s learning environment, the decisions made in these studies are motivated by higher level processes and are unlikely to be driving the memory enhancement we see here. Therefore individuals in these studies may be feeling a sense of agency, in that choosing between specific content to study requires one to be an active agent in their environment, but it is difficult to disentangle any agency-related enhancements in memory from the other higher order processes which may drive decision making.

Here, we present evidence that shows learning can be enhanced via a choice unrelated to content or timing of stimulus presentation. The participants in the current study were not making higher level decisions regarding the content of what to learn or how long to spend learning a specific item pairing. Work exploring relatively lower level volitional control has further separated active engagement in a learning environment and metacognitive decision making by giving individuals the ability to either actively or passively control the timing and content of the stimuli presentation (Markant et al., 2014, Voss 2011a, Voss 2011b). These effects could be driven by a number of processes that come along with control over learning such as increased coordination of attention, but show active control can enhance memory when not driven by higher order processes including decision making. Work from our group has established that memory can be enhanced via a choice that is unrelated to content of what is to-be-learned (Murty et al., 2015, 2019). The perceived sense of agency in these previous studies came with having a choice to flip over one of two occluder screens revealing an object. Thus the choice being made parallels the choice in the current study, where the choice made by the participant did not affect the content of the objects, which was predetermined by the experimenter, nor did it control the timing of presentation. Despite this separation between control over the learning environment and the ability to choose, this work still found memory enhancements for objects that appeared after a free choice. Further this effect was found to be independent of simple attentional manipulation as participants spent no more time viewing items on free choice compared to forced-choice trials (Murty et al., 2015). Another possibility is that individuals feel a heightened sense of ownership over items they have exerted their agency upon, such that the items become more self-relevant which has been shown to positively affect memory (Kim & Johnson, 2012). When an individual feels a high sense of agency over an item, via a choice to move an item in a certain direction, they remember that item better than items they feel a low sense of agency over (Hon & Yeo, 2021). It is possible the individuals in the current study feel a relatively higher sense of agency-on-agency trials compared to forced-choice trials, causing the item sequence to become more self-relevant. While individuals may feel a sense of self-relevance for the items that appear prior to the choice, perhaps they feel their actions are affecting that item or causing a specific outcome such that on a given agency trial the participant feels their actions caused the contestant to win the prize.

The explanations detailed above may account for enhanced memory on an item level but do not address how participants are encoding the items into an integrated sequence. Agency is possibly modulating memory to integrate items within an event into an integrated unified representation. If this were the case, individuals would use the overlapping information they are presented within the sequence to bind them together in memory. For example, in order to remember a specific contestant (A) - prize (C) pair, they would need to remember and bind the intervening pairs: contestant (A) - door (B) and door (B) - prize (C). While the individuals in the present study viewed the A-C and A-B pairs on the screen during the encoding phase, they never saw the B-C pair together. Thus, in order to integrate the sequence into a unified representation, they would not only need to recall the studied pairs (e.g. A-B) but would also need to infer the relationship of the items not seen together (e.g. B-C) which is accomplished by using the overlapping information from the other pairs to bind all three item pairs together (Shohamy & Wagner, 2008, Zeithamova & Preston, 2010, Zeithamova et al., 2012). From this, one might predict B-C and A-B memory drives memory for A-C given the inferential binding necessary to encode the sub-components of the decision sequence and that this effect would be seen across conditions as most paradigms exploring this effect use passive methods of learning. Interestingly, we show evidence that participants better remembered the A-C pair when they remembered both the A-B and B-C and had agency over that trial. This is in line with previous work showing how overlapping information from a single event can bind into an integrated sequence (Horner et al., 2015), particularly when the learners have the ability to control a minor aspect of their experience (Markant, 2020). This effect deviates from the transitive inference literature in that it was only seen when the individual had agency over the trial and it was independent of working memory capacity (not reported here, all ps > 0.40).

Another putative explanation for our sequence memory benefits is that the perceived ability to choose occurs at a critical point within a trial to influence associative memory and bind over what would otherwise serve as a context shift. In event memory and cognition, event boundaries chunk the continual stream of incoming information of an experience into smaller events (DuBrow & Davachi, 2013, Zacks et al, 2007). It has been shown that individuals can use spatial demarcation points, such as doors, as event boundaries to parse information that occurred prior to the boundary and information that occurred after the boundary (Radvansky & Zacks, 2017). It is possible that the door in the current study serves as a boundary on forced-choice trials, thus leading to worse memory for the A-C pair, but when given agency individuals can bridge over this boundary (i.e., carry the cognitive context over the boundary) to form the complete association of the sequence. While the current study cannot fully address this, future work will explore how the agency-related associative memory enhancements seen here can be used at contextual shifts between events to potentially negate their effect.

While our results are purely behavioral, they provide a theoretical framework to better understand the underlying neural systems. We propose that sequence integration by agency could potentially be caused by modulation of the hippocampus. Agency has been shown to employ similar neural mechanisms related to associative binding. Agency may endorse the formation of associations due to cortico-striatal interactions underlying the act of choosing (Delgado 2007, Leotti et al., 2010, Leotti & Delgado, 2011). These processes are thought to signal motivational significance and are known to modulate hippocampal activity (Lisman & Grace, 2005, Shohamy & Adcock, 2010, Shohamy & Turk-Browne, 2014), which is also an important factor in binding multiple elements of an experience together and which could be driving the findings reported here (Eichenbaum et al., 2007, Mayes et al., 2007, Squire et al., 2004). The hippocampus has also been shown to rapidly recognize anticipatory signals and predict upcoming action-outcomes based on past associations between pairs of stimuli across various modalities (Hindy et al., 2016, Kok & Turk-Browne, 2018, Kok et al. 2020). It is possible these mechanisms are involved in integrating the various aspects of the sequence when the participants are given the signal that they will have agency over a given trial.

In conclusion, our findings show enhanced memory for the items and the associations between items when an individual is given agency over the situation. We also show that agency may be facilitating the integration of the items into a sequence such that recollection of any given pair within the sequence is dependent on recollection of all other aspects of the sequence. This process may be dependent on hippocampal modulation by cortico-striatal interactions that come online during the act of choosing. However, further work involving neuroimaging will need to be done to definitively address these questions. Overall, our results add to a growing literature examining how agency over an item or sequence of items can influence memory and bolster associations between items.

